# Using MinION^™^ to characterize dog skin microbiota through full-length 16S rRNA gene sequencing approach

**DOI:** 10.1101/167015

**Authors:** Anna Cuscó, Joaquim Viñes, Sara D’Andreano, Francesca Riva, Joaquim Casellas, Armand Sánchez, Olga Francino

**Affiliations:** Vetgenomics, Ed Eureka, Parc de Recerca UAB, Barcelona, Spain; Molecular Genetics Veterinary Service (SVGM), Veterinary School, Universitat Autònoma de Barcelona, Barcelona, Spain

**Keywords:** MinION, nanopore, 3^rd^ generation sequencing, microbiota, microbiome, 16S rRNA, dog, canine, skin, inner pinna

## Abstract

The most common strategy to assess microbiota is sequencing specific hypervariable regions of 16S rRNA gene using 2^nd^ generation platforms (such as MiSeq or Ion Torrent PGM). Despite obtaining high-quality reads, many sequences fail to be classified at the genus or species levels due to their short length. This pitfall can be overcome sequencing the full-length 16S rRNA gene (1,500bp) by 3^rd^ generation sequencers.

We aimed to assess the performance of nanopore sequencing using MinION^™^ on characterizing microbiota complex samples. First set-up step was performed using a staggered mock community (HM-783D). Then, we sequenced a pool of several dog skin microbiota samples previously sequenced by Ion Torrent PGM. Sequences obtained for full-length 16S rRNA with degenerated primers retrieved increased richness estimates at high taxonomic level (Bacteria and Archaea) that were missed with short-reads. Besides, we were able to obtain taxonomic assignments down to species level, although it was not always feasible due to: i) incomplete database; ii) primer set chosen; iii) low taxonomic resolution of 16S rRNA gene within some genera; and/or iv) sequencing errors. Nanopore sequencing of the full-length 16S rRNA gene using MinION^™^ with 1D sequencing kit allowed us inferring microbiota composition of a complex microbial community to lower taxonomic levels than short-reads from 2^nd^ generation sequencers.

## Introduction

Bacteria, fungi, viruses and archaea are the main microorganisms constituting the microbiota, which is defined as the microbial communities inhabiting a specific environment (1). In humans, many efforts have been made to characterize the different body site ecosystems and their associated microbial communities, mainly at bacterial level (2,3), which are the most abundant microorganisms on the human-associated microbiota (4,5).

Studying host-associated microbiota has provided many insights on health and diseases for many different body sites (6,7). In human skin, alterations on skin microbiota have been associated to numerous cutaneous diseases, such as acne vulgaris (8,9), psoriasis (10–12), or atopic dermatitis (13–17). Not only humans, but also dogs presented, for example, altered microbiota states during atopic dermatitis disease (18–20).

The most common strategy to assess bacterial microbiota is amplifying and sequencing specific regions of 16S rRNA gene using 2^nd^ generation massive sequencing technologies (for a review see (21)). This bacterial marker gene is ubiquitously found in bacteria, and has nine hypervariable regions (V1-V9) that can be used to infer taxonomy (22).

The ability to classify sequences to the genus or species level is a function of read length, sample type, the reference database (23), and the quality of the sequence. High-quality short-reads obtained from 2^nd^ generation sequencers (250-350 bp) bias and limit the taxonomic resolution of this gene. The most common region amplified with Illumina MiSeq or Ion Torrent PGM^™^ for bacterial taxonomic classification is V4, but this region fails to amplify some significant species for skin microbiota studies, such as *Propionibacterium acnes.* So, when performing a skin microbiota study the preferred choice is V1-V2 regions, although they lack sensitivity for the genus *Bifidobacterium* and poorly amplify the phylum *Verrucomicrobia* (21). On the other hand, near full-length 16S rRNA gene sequences are required for accurate richness estimations especially at higher taxa (24), which are necessary on microbiota studies. Besides, full-length reference sequences are needed for performing phylogenetic analyses or designing lineage specific primers (23), especially in species different to human or mouse, in which previous metagenomics approaches deciphered the richness of bacterial species in the great and different variety of microbiome samples analyzed.

With the launching of 3^rd^ generation single-molecule technology sequencers, these short-length associated problems can be overcome by sequencing the full or almost full-length of 16S rRNA gene with different sets of universal primers (25). Results for full-length 16S rRNA gene have been reported for Pacific Biosciences (PacBio) platform (23,26–30). Schloss and collaborators reported the possibility of generating near full-length 16S rRNA gene sequences with error rates slightly higher, but comparable to the 2^nd^ generation platforms (0.03%) (23). The primary limitation on the PacBio platform is the accessibility to the sequencers and the cost of generating the data.

MinION^™^ sequencer of Oxford Nanopore Technologies (ONT) (https://nanoporetech.com) is a 3^rd^ generation sequencer that is portable, affordable with a small budget and offers long-read output (only limited by DNA extraction protocol). Besides, it can provide a rapid real-time and on-demand analysis very useful on clinical applications. Several studies targeting the full 16S rRNA gene have already been performed using MinION^™^ to: i) identify pure bacterial culture (31); ii) characterize artificial and already-characterized bacterial communities (mock community) (32–34); and to iii) characterize complex microbiota samples, from mouse gut (35), wastewater (31) and pleural effusion from a patient with empyema (34).

Here we aim to assess the potential of Nanopore sequencing in complex microbiota samples using the full-length 16S rRNA (1,500bp). First set-up step is performed using a staggered mock community (HM-783D). Then, we sequenced a pool of several skin microbiota samples previously sequenced by Ion Torrent PGM^™^.

## Material and methods

### Samples and DNA extraction

As simple microbial community, we used a Microbial Mock Community HM-783D kindly donated by BEI resources (http://www.beiresources.org) that contained genomic DNA from 20 bacterial strains with staggered ribosomal RNA operon counts (1,000 to 1,000,000 copies per organism per μL). This mock community allowed us to perform the MinION^™^ sequencing and analysis protocol set-up.

As complex microbial community, we used a sample pool from inner pinna skin microbiota of healthy dogs, which had been previously characterized using Ion Torrent PGM^™^. Skin microbiota samples were collected using Sterile Catch-All^™^ Sample Collection Swabs (Epicentre Biotechnologies) soaked in sterile SCF-1 solution (50 mM Tris buffer (pH = 8), 1 mM EDTA, and 0.5% Tween-20). Bacterial DNA was extracted from the swabs using the PowerSoil^™^ DNA isolation kit (MO BIO) (for further details on sample collection and DNA extraction see (36)).

### MinION^™^: PCR amplification and barcoding

To prepare the DNA and the library we followed the Oxford Nanopore protocol 1D PCR barcoding amplicons (SQK-LSK108), however we used the Phusion Taq polymerase rather than the LongAmp Taq recommended in this protocol. Specifically, we amplified ~1,500bp fragments of the full 16S rRNA gene.

Bacterial DNA was amplified using a nested PCR with a first round to add the 16S rRNA gene primer sets and a second round to add the barcodes. In this study we used two sets of 16S universal primers. On one hand, primer set 27F-1391R (also named S-D-Bact-0008-c-S-20 and S-D-Bact-1391-a-A-17 (37)) amplified V1-V8 hypervariable regions of 16S rRNA gene. On the other hand, primer set 27F-1492R (also named S-D-Bact-0008-c-S-20 and S-D-Bact-1492-a-A-22 (37)) amplified V1-V9 hypervariable regions of 16S rRNA gene. These two sets of universal primers are the most commonly used when assessing full-length 16S rRNA gene, because they have shown a really low non-coverage rate, even at phylum level (38). The primers used in this study are listed in Table 1 and contain some ambiguous bases previously described to make the primers more universal (25).

**Table 1.**
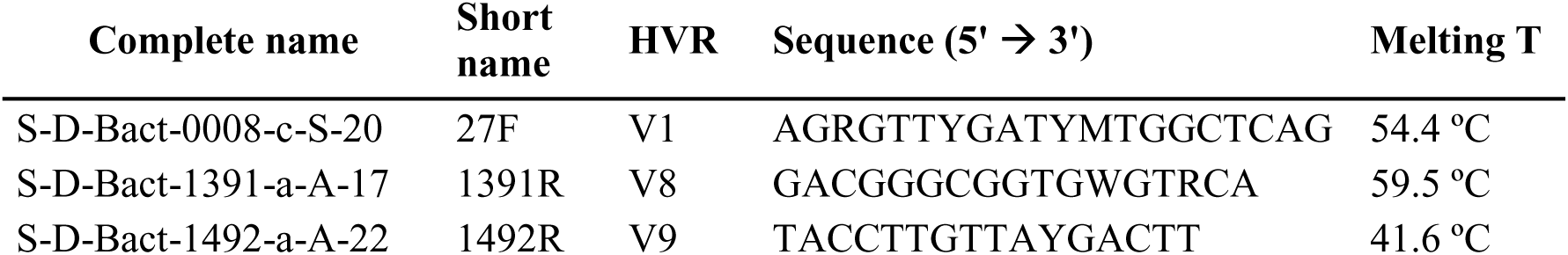
Primer sequences and hypervariable regions (HVR) targeted for full-length 16S rRNA gene amplification and sequencing.

We will distinguish among primer sets used referring to the hypervariable regions they are amplifying, so: 27F-1391R will be V1-V8; and 27F-1492R will be V1-V9.

We ordered the 16S rRNA gene primers with the Oxford Nanopore Universal Tag added to their 5’ end. The universal tag was 5’-TTTCTGTTGGTGCTGATATTGC-3’ for forward primers and 5’-ACTTGCCTGTCGCTCTATCTTC-3’ for reverse primers. These universal tags will allow the second barcoding PCR using the PCR Barcoding kit (EXP-PBC001).

In the first round, PCR mixture (25 μl) contained initial DNA sample (1 μl of DNA of the mock community and 5 μl of DNA of the skin microbiota), 5 μl of 5X Phusion Buffer HF, 0.2 mM of dNTPs, 0.02 U/μl of Phusion High Fidelity Taq Polymerase (Thermo Scientific). Primer concentrations were adapted to each primer set: for 27F-1391R, 0.4 μM of each primer and for 27F-1492R, 0.4μM of 27F and 0.8μM of 1492R. The PCR thermal profile consisted of an initial denaturation step for 30s at 98°C, followed by 25 cycles for 15s at 98°C, 15s at primer-adjusted annealing temperature, 45s at 72°C for extension, and a final step for 7 min at 72°C. The annealing temperature was adjusted to the primer set: 55ºC for 27F-1391R and 51ºC for 27F-1492R. To assess possible reagent contamination, each PCR reaction included a no template control (NTC) sample, which did not amplify.

In the second round, PCR mixture (100 μl) contained 0.5 nM of the first-round PCR product, 20 μl of 5X Phusion Buffer HF, 0.2 mM of dNTPs, 0.02 U/μl of Phusion High Fidelity Taq Polymerase (Thermo Scientific), and 2 μl of each specific barcode (EXP-PBC001) as recommended in the Oxford Nanopore protocol 1D PCR barcoding amplicons (SQK-LSK108). The PCR thermal profile consisted of an initial denaturation step for 30s at 98°C, followed by 15 cycles for 15s at 98°C, 15s at 62ºC for annealing, 45s at 72°C for extension, and a final extension step for 7 min at 72°C.

Following each PCR round, a clean-up step using AMPure XP beads at 0.5X concentration was used to discard short fragments as recommended by the manufacturer. DNA quantity was assessed using Qubit fluorimeter.

A final equimolar pool containing 1ug of the barcoded DNA samples in 45 uL of DNAse and RNAse free water were used to prepare the sequencing library.

### MinION^™^: Library preparation and sequencing

The Ligation Sequencing Kit 1D (SQK-LSK108) was used to prepare the amplicon library to load into the MinION^™^ following the instructions of the 1D PCR barcoding amplicon protocol of ONT. Input DNA samples were 1 μg of the barcoded DNA pool in a volume of 45 μL and 5 μL of DNA CS (DNA from lambda phage, used as a sequencing positive control). The DNA was processed for end repair and dA-tailing using the NEBNext End Repair / dA-tailing Module (New England Biolabs). A purification step using Agencourt AMPure XP beads (Beckman Coulter) was performed and approximately the expected 700 ng of total DNA were recovered as assessed by Qubit quantification.

For the adapter ligation step, a total of 0.2 pmol of the end-prepped DNA (approximately 200 ng of our 1,500 bp fragment) were added in a mix containing 50 μL of Blunt/TA ligase master mix (New England Biolabs) and 20 μL of adapter mix, and were incubated at room temperature for 10 min. We performed a purification step using Agencourt AMPure XP beads (Beckman Coulter) and Adapter Bead Binding buffer provided on SQK-LSK108 kit to finally obtain the DNA library.

We prepared the pre-sequencing mix (12 μL of DNA library) to be loaded by mixing it with Library Loading beads (25.5 μL) and Running Buffer with fuel mix (37.5 μL).

We used SpotON Flow Cell Mk I (R9.4) (FLO-MIN106). After the quality control, we primed the flowcell with a mixture of Running Buffer with fuel mix (RBF from SQK-LSK108) and Nuclease-free water (500 μL + 500 μL). Immediately after priming, the nanopore sequencing library was loaded in a dropwise fashion using the spot-on port.

Once the library was loaded we initiated a standard 48h sequencing protocol using the MinKNOW^™^ software.

### MinION^™^: Data pre-processing and analysis

The first flow cell contained two technical replicates of the mock community (M1 and M2) amplified with V1-V9 primer set together with other skin microbiota samples not included in this study. Basecalling was performed using the Metrichor^™^ agent 1D barcoding for pre-existing basecalls and demultiplexing using EPI2ME debarcoding workflow. Finally, *fast5* files were converted to fastq files using poRe (39) and adapters were trimmed using Porechop (40) (Figure 1).

**Figure 1.**
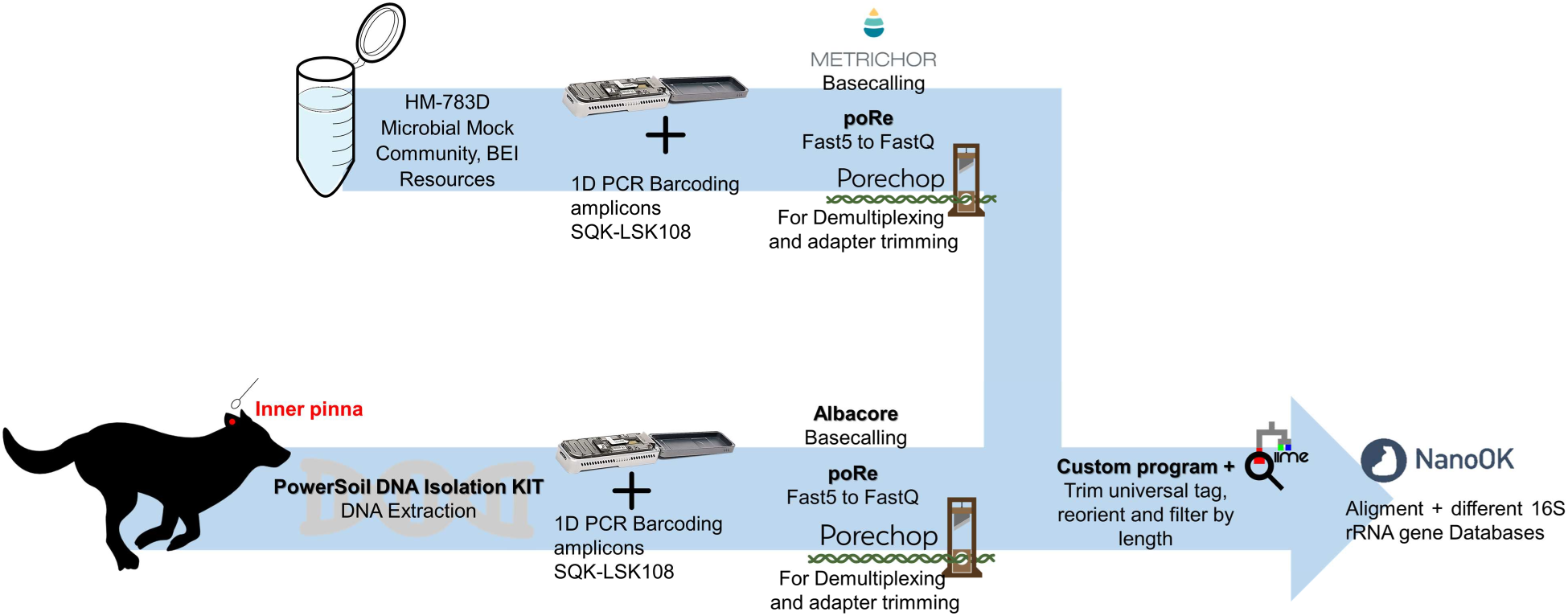
Analysis workflow of this study. HM783D was a mock community representing a simple microbial community, which was sequenced per duplicate using V1-V9 primer set (technical replicates). 5 samples from the canine inner pinna were representing a complex microbial community, and were sequenced with V1-V8 and V1-V9 (biological replicates). The same sequencing kit was used, and different pieces of software were applied in the different steps.

The second flow cell included a pool of 5 canine skin microbiota samples from the inner pinna amplified by V1-V8 and V1-V9 primer sets (biological replicates). Basecalling was performed using Albacore v0.8.4 software. Again, *fast5* files were converted to *fastq* files using poRe (39). Afterwards sequences were demultiplexed and adapters trimmed using Porechop (40) (Figure 1).

As a final step, we trimmed the universal tags of the sequences using a custom script and filtered out those sequences shorter than 1,100 bp for V1-V8 and 1,200 bp for V1-V9 amplifications respectively, using split_libraries.py from QIIME software (41).

We performed the analysis using NanoOK (42) with LAST aligner (43) against two databases: a subset of Greengenes database (44,45) and the *rrn* database (33).

– Greengenes database offers annotated, chimera-checked, and full-length 16S rRNA gene sequences and it is one of the most commonly used databases when performing microbiota analyses (44,45). We used the Greengenes database clustered at 99% of similarity to reduce redundancy and filtered out those sequences that did not reach species level or that did not have a minimum length of 1,400bp. This adapted database contained 20,745 sequences belonging to 3,147 different species.
– *rrn* database is a custom database created by Benitez-Paez and Sanz to analyze the rrn operons (33). It contains information of the ribosomal RNA operon (16S-ITS-23S genes) retrieved from bacterial genomes of GenBank at NCBI. This database contained 22,351 sequences belonging to 2,384 different species.

### IonTorrent PGM®: PCR amplification, massive sequencing and downstream analyses

V1–V2 regions of 16S rRNA gene were amplified using the widely used primer pair F27 (5′-AGAGTTTGATCCTGGCTCAG-3′) and R338 (5′-TGCTGCCTCCCGTAGGAGT-3′). PCR mixture (25 μl) contained 2 μl of DNA template, 5 μl of 5x Phusion High Fidelity Buffer, 2.5 μL of dNTPs (2 mM), 0.2 μM of each primer and 0.5 U of Phusion Hot Start II Taq Polymerase (Thermo Fisher).

The PCR thermal profile consisted of an initial denaturation of 30 sec at 98 °C, followed by 30 cycles of 15 sec at 98 °C, 15 sec at 55 °C, 20 sec at 72 °C and a final step of 7 min at 72 °C. To assess possible reagent contamination, each PCR reaction included a NTC sample.

For each amplicon, quality and quantity were assessed using Agilent Bioanalyzer 2100 and Qubit^™^ fluorometer, respectively. Both primers included sequencing adaptors at the 5′ end and forward primers were tagged with different barcodes to pool samples in the same sequencing reaction.

The 5 samples included in this study were sequenced in a pool with other skin microbiota samples. A sequencing pool included forty barcoded samples that were sequenced on an Ion Torrent^™^ Personal Genome Machine (PGM) with the Ion 318 Chip Kit v2 and the Ion PGM Sequencing 400 Kit (Life Technologies) under manufacturer’s conditions.

We will distinguish among primer sets referring the hypervariable regions they are amplifying, so: 27F-338R will be named V1-V2.

Raw sequencing reads were demultiplexed and quality-filtered using QIIME 1.9.1 (41). Reads included presented: a length greater than 300 bp; a mean quality score above 25 in sliding window of 50 nucleotides; no mismatches on the primer; and default values for other quality parameters. After that, quality-filtered reads were processed using vsearch v1.1 pipeline (46): a first de-replication step was applied, followed by clustering into operational taxonomic units (OTUs) at 97% similarity with a de *novo* approach and finally chimera checking was performed using uchime de *novo*. The raw OTU table was transferred into QIIME 1.9.1 and taxonomic assignment of representative OTUs was performed using the Ribosomal Database Project (RDP) Classifier (47) against Greengenes v13.8 database (44). Alignment of sequences was performed using PyNast (48). We sequentially applied some extra filtering steps in aligned and taxonomy-assigned OTU table to filter out: 1) sequences that belonged to Chloroplasts class; and 2) sequences representing less than 0.005% of total OTUs (as previously done in (49)). The final OTU table for skin microbiota samples can be found in Additional File 1.

## Results and discussion

### Mock community analyses

We amplified full-length 16S rRNA sequences from the staggered community with primers V1-V9 by duplicate (M1 and M2). We processed a total of 11,284 sequences for M1 and 22,995 for M2. The taxonomic results obtained are shown in Table 2 and 3, both for Greengenes and *rrn* databases.

**Table 2.** Taxonomic assignment of the mock community at genus level. Results obtained at the genus level after MinION^™^ sequencing of two replicates of the staggered mock community (M1 and M2) aligned against Greengenes and rrn databases.

| Genus | Expected abund. | Greengenes |  | *rrn* |  |
| --- | --- | --- | --- | --- | --- |
|  |  | % of reads M1 | % of reads M2 | % of reads M1 | % of reads M2 |
| *Streptococcus* | ++++ | 33.99 | 33.44 | 35.44 | 34.82 |
| *Staphylococcus* | ++++ | 24.00 | 24.30 | 24.32 | 24.73 |
| *Escherichia* | ++++ | 15.41 | 16.52 | 22.91 | 24.23 |
| *Rhodobacter* | ++++ | 8.58 | 7.21 | 8.35 | 6.99 |
| *Bacillus* | +++ | 3.66 | 3.69 | 3.28 | 3.37 |
| *Clostridium* | +++ | 1.31 | 1.46 | 1.25 | 1.3 |
| *Pseudomonas* | +++ | 1.04 | 1.05 | 1.07 | 1.01 |
| *Listeria* | ++ | 0.27 | 0.30 | 0.20 | 0.30 |
| *Neisseria* | ++ | 0.17 | 0.27 | 0.17 | 0.30 |
| *Propionibacterium* | ++ | 0.17 | 0.22 | 0.13 | 0.17 |
| *Lactobacillus* | ++ | 0.27 | 0.17 | 0.28 | 0.18 |
| *Helicobacter* | ++ | 0.10 | 0.14 | 0.12 | 0.13 |
| *Acinetobacter* | ++ | 0.13 | 0.12 | 0.13 | 0.11 |
| *Enterococcus* | + | 0.14 | 0.11 | 0.12 | 0.06 |
| *Bacteroides* | + | 0.02 | 0.01 | 0.02 | 0.01 |
| *Deinococcus* | + | 0.02 | 0 | 0.02 | 0.01 |
| *Actinomyces* | + | 0 | 0 | 0 | 0 |
| Other genera* | - | 10.72 | 10.99 | 2.19 | 2.28 |

**Table 3.** Taxonomic assignment of the mock community at species level. Results obtained after MinION^™^ sequencing of two replicates of the staggered mock community (M1 and M2) aligned against Greengenes and rrn databases. Relative abundances that correlated to operon counts are in bold. (*) Is the species annotated in the database? (n.d.): not detected

| Taxonomy | Greengenes database |  |  |  | rrn database |  |  |
| --- | --- | --- | --- | --- | --- | --- | --- |
|  | n° of operons | Tax at sps?* | % M1 | % M2 | Tax at sps?* | % M1 | % M2 |
| *Escherichia coli* | 1,000,000 | Yes | 15.29 | 16.45 | Yes | 22.49 | 23.92 |
| *Rhodobacter sphaeroides* | 1,000,000 | Yes | 8.54 | 7.19 | Yes | 8.18 | 6.85 |
| *Staphylococcus epidermidis* | 1,000,000 | Yes | 18.81 | 19.04 | Yes | 15.99 | 16.71 |
| *Streptococcus mutans* | 1,000,000 | No | - | - | Yes | 17.34 | 17.12 |
| *Bacillus cereus* | 100,000 | Yes | 2.26 | 2.1 | Yes | 0.93 | 0.96 |
| *Clostridium beijerinckii* | 100,000 | No | - | - | Yes | 0.46 | 0.67 |
| *Pseudomonas aeruginosa* | 100,000 | Yes | 0.01 | 0.03 | Yes | 0.85 | 0.88 |
| *Staphylococcus aureus* | 100,000 | Yes | 1.59 | 1.81 | Yes | 6.81 | 6.49 |
| *Streptococcus agalactiae* | 100,000 | Yes | 1.27 | 1.42 | Yes | 2.36 | 2.36 |
| *Acinetobacter baumannii* | 10,000 | No | - | - | Yes | 0.11 | 0.09 |
| *Helicobacter pylori* | 10,000 | Yes | 0.1 | 0.14 | Yes | 0.11 | 0.13 |
| *Lactobacillus gasseri* | 10,000 | No | - | - | Yes | 0.17 | 0.12 |
| *Listeria monocytogenes* | 10,000 | Yes | 0.21 | 0.29 | Yes | 0.12 | 0.23 |
| *Neisseria meningitidis* | 10,000 | No | - | - | Yes | 0.14 | 0.24 |
| *Propionibacterium acnes* | 10,000 | Yes | 0.17 | 0.22 | Yes | 0.03 | 0.05 |
| *Actinomyces odontolyticus* | 1,000 | Yes | n.d. | n.d. | Yes | n.d. | n.d. |
| *Bacteroides vulgatus* | 1,000 | No | - | - | Yes | 0.02 | 0.01 |
| *Deinococcus radiodurans* | 1,000 | No | - | - | No | - | - |
| *Enterococcus faecalis* | 1,000 | No | - | - | Yes | 0.11 | 0.02 |
| *Streptococcus pneumoniae* | 1,000 | No | - | - | Yes | 3.63 | 3.38 |

Greengenes is one of the most commonly used databases in microbiota studies because it is curated and checked for chimeras (45). We used a subset of this database, which contained 20,745 sequences belonging to 3,147 different species. However, only 11 out of the 20 bacterial species of the mock community were annotated in the database down to the species level, so we expected seeing only genus level for those specific taxa (marked as “No” in Table 2).

*rrn* database (33) contains information of the ribosomal RNA operon (16S-ITS-23S genes) for 22,351 sequences belonging to 2,384 different species. This database lacks information for only one member of the mock community (*Deinococcus radiodurans*).

At the genus level, we were able to identify all the members of the mock community except *Actinomyces* even when it was represented on both databases (Table 2). The overall trend is that relative abundances correlated to operon counts: taxa with >10% of relative abundance represent operons with 1,000,000 copies; taxa with >1% of relative abundance represent operons with 100,000 copies; and successively. The exceptions were *Actinomyces* and *Rhodobacter* with lower abundances than expected that suggesting primer set V1-V9 was not optimal for these bacterial genera.

Approximately 10% of the total reads for Greengenes and 2% for *rrn* database belonged to other genera theoretically not present in the mock community. Among those “other genera”, the most abundant belonged to *Shigella, Enterobacter* and *Salmonella* that present a 16S rRNA gene with high similarity to *Escherichia coli* (present in the mock community) (50). If we consider these taxa were probably wrongly assigned because they are closely related to *Escherichia coli*, only ~2.5% and ~0.5 % of the reads aligned to Greengenes and *rrn* database respectively belong to unexpected other genera, which could be either due to sequencing errors and wrong taxonomical assignation or to cross-contamination from dog skin microbiota samples.

Delving to species level, Greengenes contains taxonomic annotation for 11 out of 20 bacterial species included in the mock bacterial community. From these, we were able to detect all of them (with the exception of *Actinomyces odontolyticus*). *Rhodobacter sphaeroides, Pseudomonas aeruginosa* were detected at lower abundances than expected. Moreover, it’s worthy to note that despite *Streptococcus mutans* was not in Greengenes database at species level, we detected the closely related species *Streptococcus sobrinus* in high abundance (M1=14.4% and M2=11.3%); in fact both belong to the *mutans* group (51). Moreover, we saw *Streptococcus infantis* (M1=9.6 and M2= 7.9%) as another abundant species. On the other hand, *rrn* database contained species level information from 19 out of the 20 bacterial species of the mock community and we were able to detect all of them (with the exception of *Actinomyces odontolyticus*) (Table 3).

When looking at the results of *rrn* database, we could see that not only *Rhodobacter sphaeroides* and *Pseudomonas aeruginosa* were underrepresented but also *Bacillus cereus* and *Clostridium beijerinckii*. Finally, *Streptococcus pneumoniae* was overrepresented, probably suggesting that the sequencing errors together with the large amount of *Streptococcus* entries in the *rrn* database (44 *Streptococcus* species in *rrn* vs 9 *Streptococcus* species in Greengenes) produced an incorrect identification of this species.

We can conclude from mock community analyses that full-length 16S rRNA sequencing with MinION^™^ is able to detect taxonomy assignments and retrieve diversity information, provided that the target species are in the database. At the genus level, we were able to accurately retrieve the mock community composition. It’s also worthy to note the good technical replicates obtained for M1 and M2 samples. Some of the biases observed in the expected taxonomic profile could be due to: i) sequencing errors; ii) primer set used; iii) low taxonomic resolution of 16S rRNA gene within some genera; and/or iv) incomplete database.

### Evaluation of primer sets V1-V8 and V1-V9 in microbial richness

Dog skin microbiota samples were sequenced as a pool with MinION^™^ after amplification of full-length 16S rRNA gene with primers targeting regions V1-V8 or V1-V9 (see Table 1). These complex microbiota samples were basecalled with Albacore v0.8.4 and *fast5* files were converted to *fastq* files. After demultiplexing, adapters and universal tags were trimmed and sequences analyzed using NanoOK (42) with LAST aligner (43). We finally obtained a total of 79,083 sequences for V1-V9 and 74,243 for V1-V8.

The same samples had been previously sequenced individually with Ion Torrent PGM^™^ with primers targeting V1-V2 hypervariable regions. In that case sequences were analyzed with QIIME 1.9.1 (41) with operational taxonomic units (OTUs) picking a representative sequence of a group of sequences with a 97% similarity and taxonomy was assigned with RDP classifier (47) against the whole Greengenes database (it contains many entries that do not reach low taxonomic levels) (44,45). Using RDP classifier, if the taxonomy assignment does not reach a specific threshold, the sequences included in the OTU are set as “Other”. We finally obtained a total of 249,572 sequences for V1-V2 region (Additional File 1).

We performed the evaluation of the primer sets using exclusively the Greengenes database, because V1-V2 short-reads were analyzed using this database. We used a subset of the Greengenes database that contained only those taxa that reached species level for the long-reads obtained for V1-V8 and V1-V9 regions with MinION^™^. We compared diversity estimates of higher taxa (from kingdom to order).

Both V1-V8 and V1-V9 primer sets for long-reads were able to retrieve more bacterial taxa than V1-V2 short reads. The bacterial richness was higher at different taxonomic levels when assessed with long-reads rather than with short-reads, as seen in Table 4, and this trend increased as we were lowering taxonomic level.

At the highest taxonomic level, we were able to detect not only Bacteria but also Archaea kingdom, despite using universal primers specific for Bacteria (25). However, they were present at really low proportions (< 0.01% of total reads), which agrees with previous results on human skin microbiota samples (52).

**Table 4.** Bacterial richness estimates for skin microbiota samples. Bacterial richness retrieved with different primer pairs targeting short (V1-V2) or full-length (V1-V8 and V1-V9) 16S rRNA gene.

|  | **V1-V2**<br>**(~350 bp)** | **V1-V8**<br>**(~1300 bp)** | **V1-V9**<br>**(~1400 bp)** |
| --- | --- | --- | --- |
| Kingdom | 1 | 2 | 2 |
| Phylum | 16 | 22 | 22 |
| Class | 33 | 46 | 47 |
| Order | 53 | 88 | 91 |

Delving into phylum level, we detected that the most common and better characterized phyla were retrieved by all the primers. These taxa represented >98% of the total skin microbiota composition. Long-read primers were able to detect 8 phyla previously unseen using V1-V2 short-reads (Table 5). It has already been reported the low coverage of this primer set for some specific phyla (21,53). Some phyla were only detected with a specific primer set, such as Lentisphaerae with V1-V8 or Fibrobacteres with V1-V9. On the other hand, GN02, TM7 and Thermi phyla belong to candidate divisions and none of their members have been cultivated (54), so databases do not have taxonomy information down to species level. Thus, since we used the Greengenes subset database with species-level sequences, no representative of those phyla were included for taxonomy assignment of V1-V8 and V1-V9 long-reads and that is probably the reason why they are only detected with V1-V2 primers.

**Table 5.** Microbial richness on skin samples. Table containing all the observed phyla per primer subset. Long-reads taxonomy was obtained from a species-level subset of Greengenes database (see materials and methods) that did not contain information of GN02, SR1 and TM7. *phyla that belong to Archaea kingdom. ** phyla with no representative at the species level in the database used for assigning taxonomy to long-reads (V1-V8 and V1-V9).

| Phylum | Long-reads |  | Short-reads |
| --- | --- | --- | --- |
|  | V1-V9 | V1-V8 | V1-V2 |
| [Thermi] | + | + | + |
| Acidobacteria | + | + | + |
| Actinobacteria | + | + | + |
| Bacteroidetes | + | + | + |
| Chlorobi | + | + | + |
| Chloroflexi | + | + | + |
| Cyanobacteria | + | + | + |
| Deferribacteres | + | + | + |
| Firmicutes | + | + | + |
| Fusobacteria | + | + | + |
| Proteobacteria | + | + | + |
| Spirochaetes | + | + | + |
| Tenericutes | + | + | + |
| Aquificae | + | + |  |
| Chlamydiae | + | + |  |
| Crenarchaeota* | + | + |  |
| Elusimicrobia | + | + |  |
| Euryarchaeota* | + | + |  |
| Planctomycetes | + | + |  |
| Synergistetes | + | + |  |
| Verrucomicrobia | + | + |  |
| Fibrobacteres | + |  |  |
| Lentisphaerae |  | + |  |
| GN02** |  |  | + |
| SR1** |  |  | + |
| TM7** |  |  | + |
| **Total** | **22/26** | **22/26** | **16/26** |

So, we were able to detect previously unseen bacteria phyla on dog skin using MinION^™^ long-amplicons for full-length 16S rRNA sequences, which provided better richness estimates. We cannot discard that this increased richness could also be due to the primers used for long amplicons, which presented some degenerated positions. Although these previously unseen bacteria phyla presented low relative abundances on canine skin microbiota, the use of long-amplicons in more uncharacterized environments will provide better diversity estimates.

### Skin microbiota analyses

We assessed canine skin microbiota composition using MinION^™^ and full-length 16S rRNA gene in a pool of 5 inner pinna samples that were previously individually sequenced with Ion Torrent PGM^®^ using V1-V2 short-reads. In Figure 2a we can see the microbiota profile of the most abundant taxa (> 0.5% of total relative abundance) per individual sample included in the pool when sequencing V1-V2 region with Ion Torrent PGM®. We can see that all of them have been identified down to family level and some of them also to genus level.

**Figure 2.**
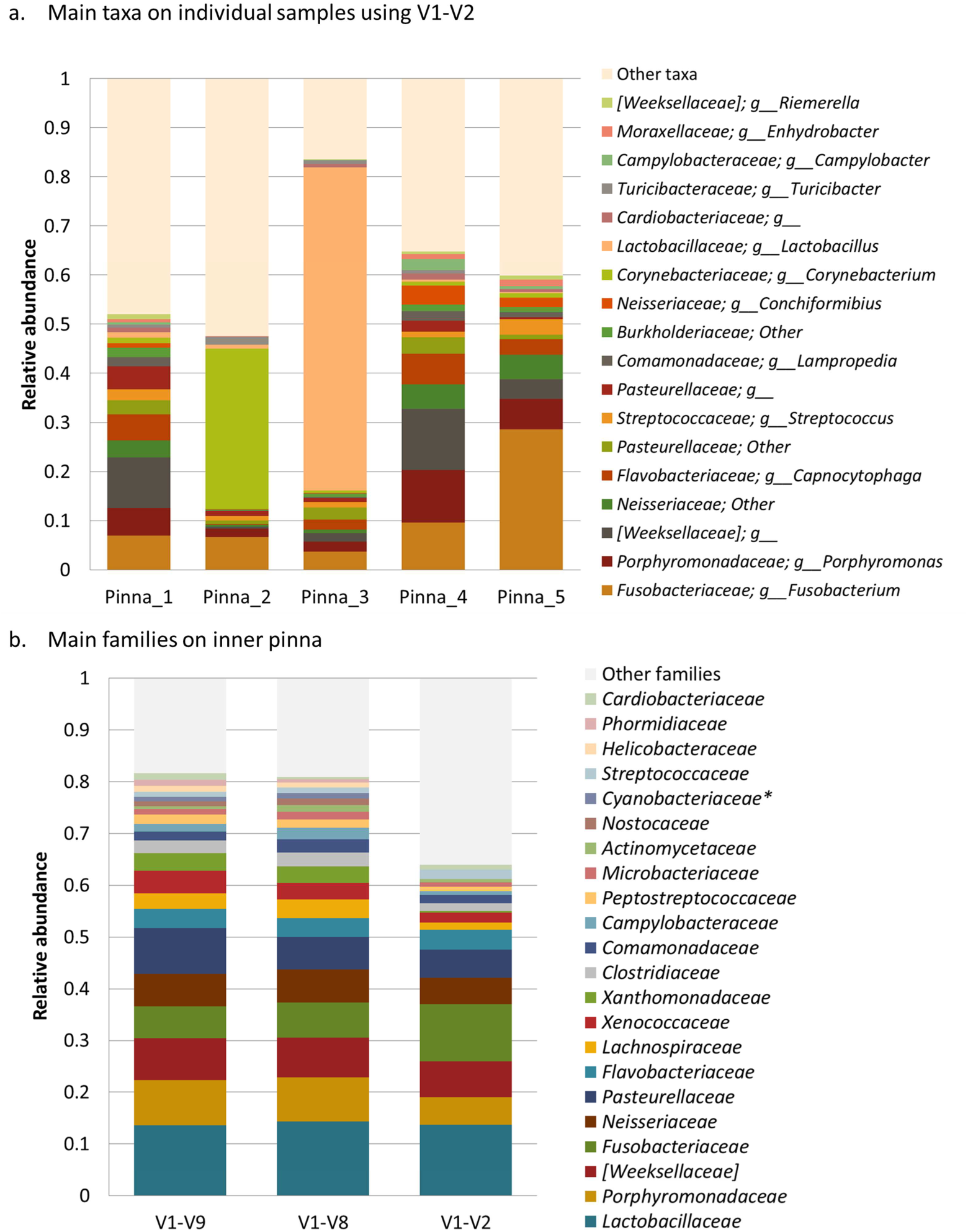
Skin microbiota taxonomic profile using long and short reads. Only abundant taxa included (>0.5% of total relative abundance). Bar plots of (a) the most abundant taxa on individual skin samples sequenced with V1-V2 primers in short-reads. “g__” means there is no information at genus level; and b) the main families on dog skin microbiota when sequencing the full-length 16S rRNA gene (V1-V9 and V1-V8) and when averaging the V1-V2 individual results.

We have averaged the results from V1-V2 regions (sequenced by IonTorrent PGM®) to compare the taxonomic profile with that obtained from V1-V8 and V1-V9 regions (sequenced by MinION^™^) (Figure 2b). The global taxonomic profile at family level was equivalent when comparing V1-V8 and V1-V9 that are biological replicates. V1-V2 taxonomic profile was similar for the abundant species, and differed when looking at low-abundant taxa. The pools used for V1-V8 and V1-V9 nanopore sequencing were not equally representing all the individual samples despite working with an equimolar pool. Moreover, *Lactobacillaceae* seems to be an abundant family on dog inner’s pinna and in fact is almost exclusive to a unique sample (18A).

The pool of canine inner pinna samples was sequenced twice using a different set of primers (V1-V8 and V1-V9) and gave highly similar taxonomic results within each database, making the taxonomic assignment robust (Figure 3 and Additional File 2). However, mock community results showed that species-level resolution was challenging even with full-length 16S rRNA because sometimes the target species was not present in the database and the assigned taxonomy corresponded to a closed-related species rather than the actual one. Because of each database contained different taxonomic annotations, we considered a species-level assignment reliable when the two unrelated databases gave identical annotation.

**Figure 3.**
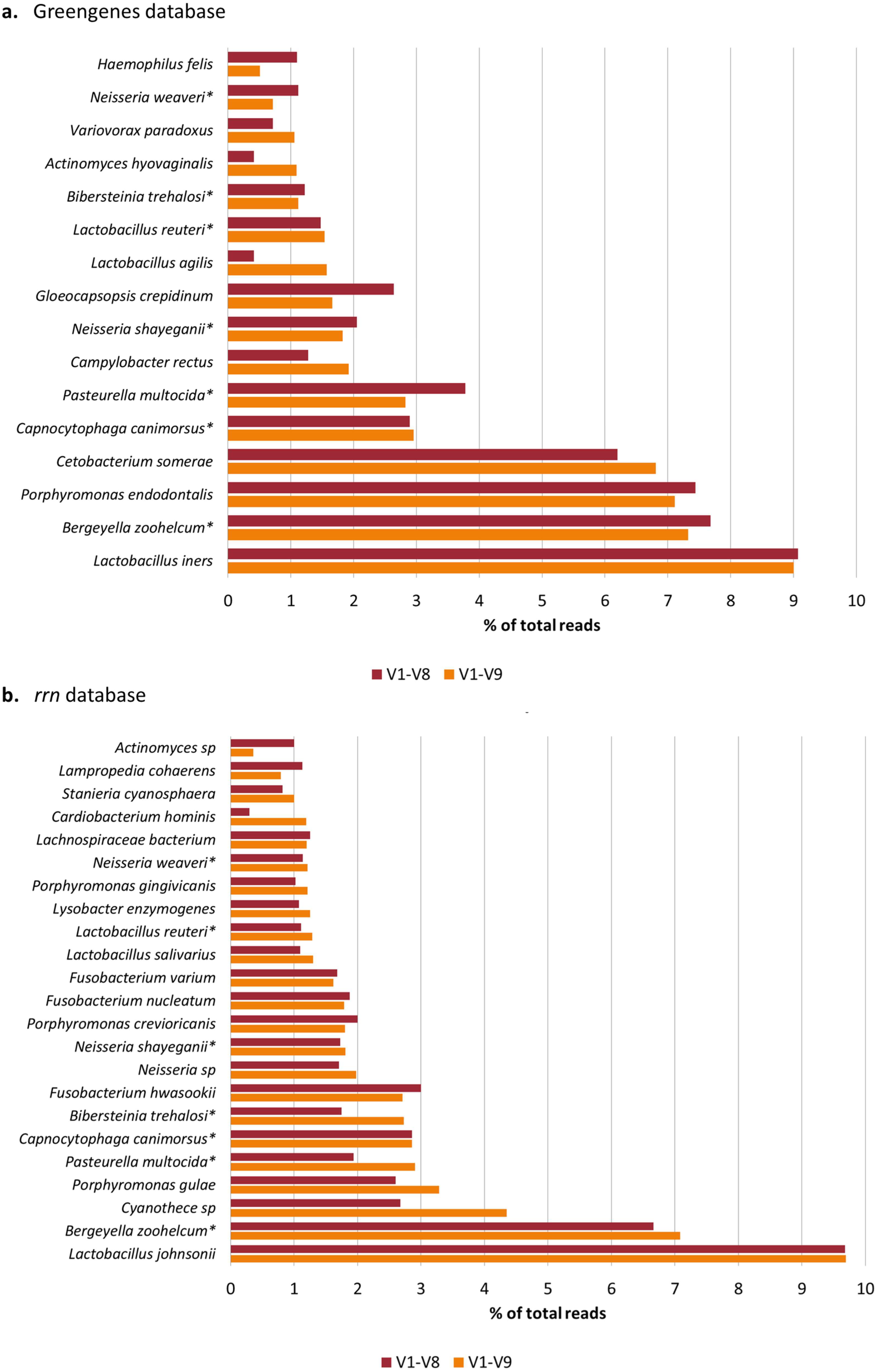
Skin microbiota composition at species level. Comparison of the abundant species (>1% of the total microbiota composition) detected against the **(a)** Greengenes and **(b)** *rrn* databases in the dog skin microbiota samples amplified with V1-V8 and V1-V9 16S rRNA primers and sequenced with MinION^™^. Additional File 2 contains all the taxa identified (*) coincident taxonomic assignments using independent databases.

When comparing the most abundant species (>1% of total relative abundances), we could see 7 bacterial species identified by two independent databases: *Bergeyella zoohelcum, Capnocytophaga canimorsus, Pasteurella multocida, Neisseria shayeganii, Lactobacillus reuteri, Bibersteinia trehalosi*, and *Neisseria weaver* (Figure 3). Other 13 bacterial species presented the same identical taxonomy assignment in the two databases with lower abundances (>0.01%) (Table 6).

**Table 6.** Bacterial species identified at canine skin microbiota confirmed by Greengenes and rrn databases. % of total skin microbiota composition.

| Species matching GG and rrn databases (%) | GG |  | rrn |  |
| --- | --- | --- | --- | --- |
|  | V1-V8 | V1-V9 | V1-V8 | V1-V9 |
| *Bergeyella zoohelcum* | 7.33 | 7.68 | 6.71 | 7.13 |
| *Porphyromonas endodontalis* | 7.12 | 7.44 | 0.32 | 0.19 |
| *Pasteurella multocida* | 2.83 | 3.78 | 1.94 | 2.97 |
| *Capnocytophaga canimorsus* | 2.96 | 2.89 | 2.87 | 2.87 |
| *Neisseria shayegani* | 1.83 | 2.05 | 1.74 | 1.82 |
| *Lactobacillus reuteri* | 1.54 | 1.48 | 1.12 | 1.27 |
| *Bibersteinia trehalosi* | 1.12 | 1.22 | 1.76 | 2.74 |
| *Neisseria weaveri* | 0.71 | 1.12 | 1.15 | 1.22 |
| *Lactobacillus salivarius* | 0.71 | 0.91 | 1.12 | 1.31 |
| *Lactobacillus ruminis* | 0.43 | 0.67 | 0.45 | 0.56 |
| *Anabaena cylindrica* | 0.50 | 0.53 | 0.89 | 0.67 |
| *Stanieria cyanosphaera* | 0.31 | 0.47 | 0.83 | 1.00 |
| *Haemophilus parasuis* | 0.31 | 0.44 | 0.66 | 0.85 |
| *Desulfomicrobium orale* | 0.31 | 0.41 | 0.29 | 0.41 |
| *Gloeobacter violaceus* | 0.34 | 0.35 | 0.19 | 0.12 |
| *Clostridium acidurici* | 0.43 | 0.31 | 0.33 | 0.26 |
| *Pseudoxanthomonas dokdonensis* | 0.13 | 0.3 | 0.14 | 0.18 |
| *Neisseria wadsworthii* | 0.12 | 0.19 | 0.17 | 0.16 |
| *Cylindrospermum stagnale* | 0.15 | 0.11 | 0.19 | 0.08 |
| *Actinomyces europaeus* | 0.24 | 0.09 | 0.15 | 0.02 |

When comparing the taxonomic information obtained by V1-V2 (Figure 2a) to that obtained with long-reads (Figure 3) we could reach lower taxonomic levels for: 1) [*Weeksellaceae*] family, with *Bergeyella zoohelcum*; 2) *Neisseriaceae,* with *Neisseria shayeganii* and *Neisseria weaveri;* 3) *Pasteurellaceae*, with *Pasteurella multocida* and *Bibersteinia trehalosi;* and 4) *Flavobacteriaceae*, with *Capnocitophaga canimorsus.*

Some of these species detected on dog inner pinna skin microbiota have been previously isolated or detected in dogs. *Bergeyella zoohelcum* and *Neisseria shayeganii* have been detected in canine oral microbiota of healthy dogs, as well as *Pasteurellaceae sp*. and *Capnocytophaga* (55). In fact, *Bergeyella zoohelcum, Capnocytophaga canimorsus* and *Pasteurella multocida* are classically considered as zoonotic pathogens because they can be responsible for bacteremia produced after dog bites (56–58). Also *Neisseria weaveri* was associated to dog bites (59). On the other hand, *Lactobacillus reuteri* was one of the most prevalent and abundant lactic acid bacteria isolated from fecal and intestinal microbiota of healthy dogs (60,61). Thus, these taxa are likely to be normal inhabitants of the skin microbiota of healthy dogs.

We could also see that *Fusobacteriaceae* (specifically *Fusobacterium* genus) was one of the most abundant families on the inner pinna skin microbiota with V1-V2 short-reads. However, the subset of Greengenes database used lacked entries of *Fusobacterium* at the species level so this genus is not detected. *Porphyromonadaceae* was another abundant taxon of dog skin microbiota, but the species level taxonomy differed when using each database. The only representative at species level in Greengenes is *P. endodontalis,* whereas rrn database contains 9 different species. When analyzing the results against Greengenes, all the *Porphyromonas* sequences are classified as *P. endodontalis*. On the other hand when aligning the same sequences against rrn database, these are classified in 7 different *Porphyromonas* species (see rrn summary in Additional File 2). This result highlights again the need databases that contain the most relevant taxa at species level.

## Conclusions

Nanopore sequencing of the full-length 16S rRNA gene with MinION^™^ allowed us inferring microbiota composition from both simple and complex microbial communities. Moreover, long-reads and degenerated primers were able to retrieve increased richness estimates, which show us previously unseen phyla on dog skin microbiota, despite being at really low abundances.

Taxonomy assignment down to species level was obtained, although it was not always feasible due to: i) sequencing errors; ii) primer set chosen; iii) low taxonomic resolution of 16S rRNA gene within some genera; and/or iv) incomplete database.

With the nanopore reads, we assigned taxonomy through alignment strategies executing all-vs-all comparisons that need many computational resources, so we needed a small database to obtain results. When working with a 16S database subset, we should be sure to include the most relevant taxa even if they do not have representative members at species level. Oxford Nanopore Technologies offers other bioinformatics tools, such as the cloud-based EPI2ME platform. However, we run out of memory on our hard disk when trying to perform these analyses.

Future studies should be relying on the new 1D^2^ kit that presents higher accuracy and also other experimental strategies could be assessed to perform microbiota studies taking profit of 3^rd^ generation sequencing (Box 1). Other amplicon-based strategies have shown potential, such as: i) sequencing the whole rrn operon constituted by 16S rRNA-ITS-23S rRNA (33); ii) sequencing the cDNA from size selected SSU rRNA (62); or iii) sequencing other bacterial barcodes, such as cpn60 (63). Another potential approach would be sequencing the 16S rRNA directly that now is feasible with nanopore sequencing, Smith and colleagues were able to detect the modified bases of *Escherichia coli* that can identify a pathogenic strain (64). Finally, metagenomics approach would allow seeing not only taxonomic information of the whole community but also potential functions of the community (genes).

### Box 1.

**Alternative strategies to assess microbiota**

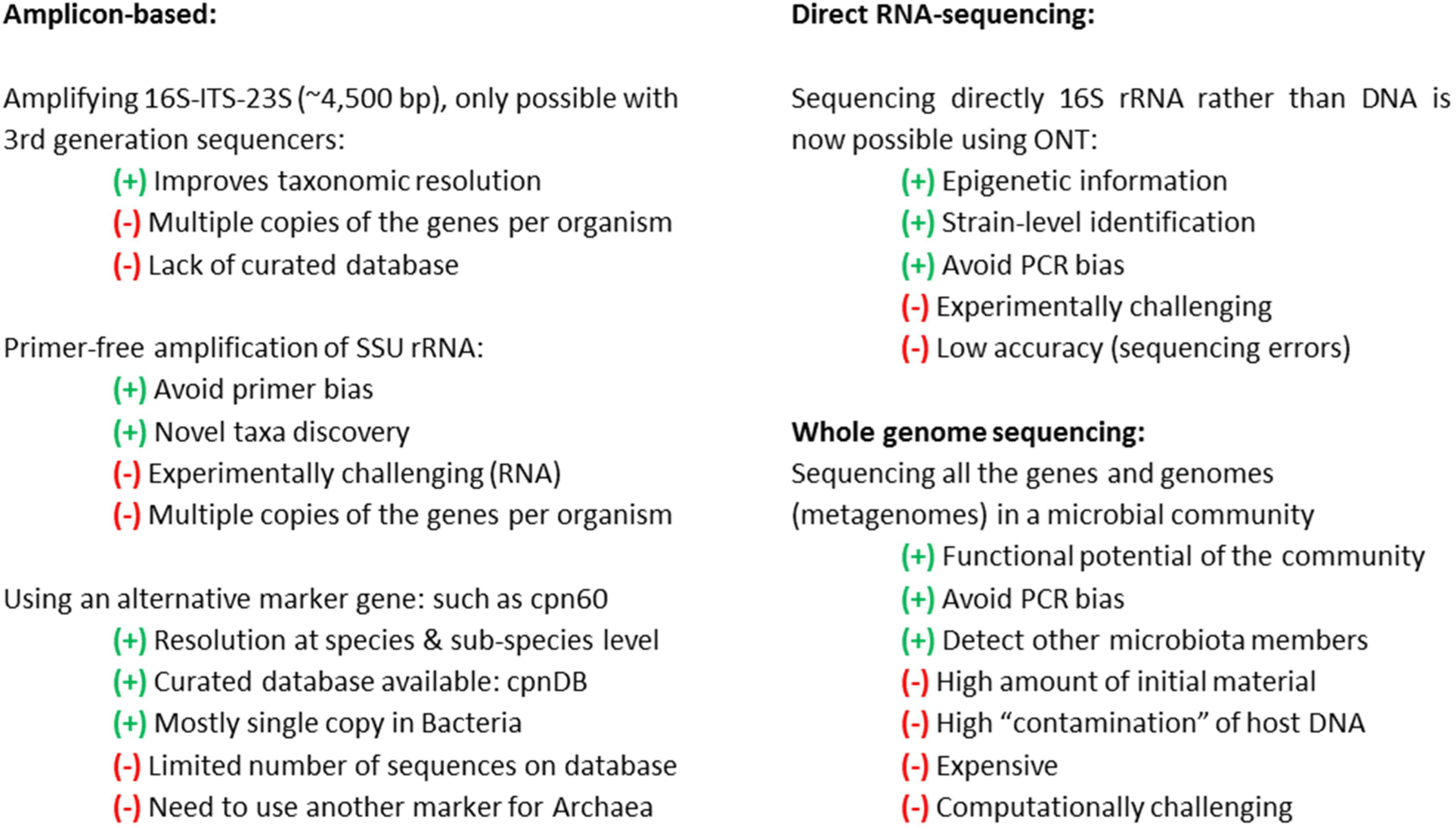

## Acknowledgments

This work was supported by a grant awarded by Generalitat de Catalunya to AC (Industrial Doctorate program, 2013 DI 011) and SD (Industrial Doctorate program, 2015 DI 044).

## References

1. Marchesi JR, Ravel J. The vocabulary of microbiome research: a proposal. Microbiome (2015) 3:31. doi:10.1186/s40168-015-0094-5

2. Costello EK, Lauber CL, Hamady M, Fierer N, Gordon JI, Knight R. Bacterial community variation in human body habitats across space and time. Science (2009) 326:1694–1697. doi:10.1126/science.1177486

3. The Human Microbiome Project Consortium. Structure, function and diversity of the healthy human microbiome. Nature (2012) 486:207–214. doi:10.1038/nature11234

4. Qin J, Li R, Raes J, Arumugam M, Burgdorf KS, Manichanh C, Nielsen T, Pons N, Levenez F, Yamada T, et al. A human gut microbial gene catalogue established by metagenomic sequencing. Nature (2010) 464:59–65. doi:10.1038/nature08821

5. Belkaid Y, Segre JA. Dialogue between skin microbiota and immunity. Science (2014) 346:954–9. doi:10.1126/science.1260144

6. Thaiss CA, Zmora N, Levy M, Elinav E. The microbiome and innate immunity. Nature (2016) 535:65–74. doi:10.1038/nature18847

7. Belkaid Y, Hand TW. Role of the microbiota in immunity and inflammation. Cell (2014) 157:121–141. doi:10.1016/j.cell.2014.03.011

8. Fitz-Gibbon S, Tomida S, Chiu BH, Nguyen L, Du C, Liu M, Elashoff D, Erfe MC, Loncaric A, Kim J, et al. *Propionibacterium acnes* strain populations in the human skin microbiome associated with acne. J Invest Dermatol (2013) 133:2152–60. doi:10.1038/jid.2013.21

9. Dreno B, Martin R, Moyal D, Henley JB, Khammari A, Seité S. Skin microbiome and acne vulgaris: *staphylococcus*, a new actor in acne. Exp Dermatol (2017) doi:10.1111/exd.13296

10. Gao Z, Tseng C, Strober BE, Pei Z, Blaser MJ. Substantial alterations of the cutaneous bacterial biota in psoriatic lesions. PLoS One (2008) 3:e2719. doi:10.1371/journal.pone.0002719

11. Fahlén A, Engstrand L, Baker BS, Powles A, Fry L. Comparison of bacterial microbiota in skin biopsies from normal and psoriatic skin. Arch Dermatol Res (2012) 304:15–22. doi:10.1007/s00403-011-1189-x

12. Alekseyenko A V, Perez-Perez GI, De Souza A, Strober B, Gao Z, Bihan M, Li K, Methé BA, Blaser MJ. Community differentiation of the cutaneous microbiota in psoriasis. Microbiome (2013) 1:31. doi:10.1186/2049-2618-1-31

13. Kong HH, Oh J, Deming C, Conlan S, Grice EA, Beatson MA, Nomicos E, Polley EC, Komarow HD, Murray PR, et al. Temporal shifts in the skin microbiome associated with disease flares and treatment in children with atopic dermatitis. Genome Res (2012). 22:850-9. doi:10.1101/gr.131029.111

14. Oh J, Freeman AF, Park M, Sokolic R, Candotti F, Holland SM, Segre JA, Kong HH. The altered landscape of the human skin microbiome in patients with primary immunodeficiencies. Genome Res (2013) 23:2103–14. doi:10.1101/gr.159467.113

15. Seite S, Flores GE, Henley JB, Martin R, Zelenkova H, Aguilar L, Fierer N. Microbiome of affected and unaffected skin of patients with atopic dermatitis before and after emollient treatment. J Drugs Dermatol (2014) 13:1365–72.

16. Chng KR, Tay ASL, Li C, Ng AHQ, Wang J, Suri BK, Matta SA, McGovern N, Janela B, Wong XF, et al. Whole metagenome profiling reveals skin microbiome-dependent susceptibility to atopic dermatitis flare. Nat Microbiol (2016) 1:16106. doi:10.1038/nmicrobiol.2016.106

17. Kennedy EA, Connolly J, Hourihane JO, Fallon PG, McLean WHI, Murray D, Jo JH, Segre JA, Kong HH, Irvine AD. Skin microbiome before development of atopic dermatitis: Early colonization with commensal staphylococci at 2 months is associated with a lower risk of atopic dermatitis at 1 year. J Allergy Clin Immunol (2017) 139:166–72. doi:10.1016/j.jaci.2016.07.029

18. Rodrigues Hoffmann A, Patterson AP, Diesel A, Lawhon SD, Ly HJ, Elkins Stephenson C, Mansell J, Steiner JM, Dowd SE, Olivry T, et al. The skin microbiome in healthy and allergic dogs. PLoS One (2014) 9:e83197. doi:10.1371/journal.pone.0083197

19. Bradley CW, Morris DO, Rankin SC, Cain CL, Misic AM, Houser T, Mauldin EA, Grice EA. Longitudinal evaluation of the skin microbiome and association with microenvironment and treatment in canine atopic dermatitis. J Invest Dermatol (2016) 136:1182-90. doi:10.1016/j.jid.2016.01.023

20. Pierezan F, Olivry T, Paps JS, Lawhon SD, Wu J, Steiner JM, Suchodolski JS, Rodrigues Hoffmann A. The skin microbiome in allergen-induced canine atopic dermatitis. Vet Dermatol (2016) 27:332-e82. doi:10.1111/vde.12366

21. Kuczynski J, Lauber CL, Walters W a., Parfrey LW, Clemente JC, Gevers D, Knight R. Experimental and analytical tools for studying the human microbiome. Nat Rev Genet (2011) 13:47–58. doi:10.1038/nrg3129

22. Clarridge JE. Impact of 16S rRNA Gene Sequence Analysis for Identification of Bacteria on Clinical Microbiology and Infectious Diseases. Clin Microbiol Rev (2004) 17:840–862. doi:10.1128/CMR.17.4.840-862.2004

23. Schloss PD, Jenior ML, Koumpouras CC, Westcott SL, Highlander SK. Sequencing 16S rRNA gene fragments using the PacBio SMRT DNA sequencing system. PeerJ (2016) 4:e1869. doi:10.7717/peerj.1869

24. Yarza P, Yilmaz P, Pruesse E, Glöckner FO, Ludwig W, Schleifer K-H, Whitman WB, Euzéby J, Amann R, Rosselló-Móra R. Uniting the classification of cultured and uncultured bacteria and archaea using 16S rRNA gene sequences. Nat Rev Microbiol (2014) 12:635–645. doi:10.1038/nrmicro3330

25. Klindworth A, Pruesse E, Schweer T, Peplies J, Quast C, Horn M, Glockner FO. Evaluation of general 16S ribosomal RNA gene PCR primers for classical and next-generation sequencing-based diversity studies. Nucleic Acids Res (2013) 41:e1. doi:10.1093/nar/gks808

26. Fichot EB, Norman RS. Microbial phylogenetic profiling with the Pacific Biosciences sequencing platform. Microbiome (2013) 1:10. doi:10.1186/2049-2618-1-10

27. Mosher JJ, Bernberg EL, Shevchenko O, Kan J, Kaplan LA. Efficacy of a 3rd generation high-throughput sequencing platform for analyses of 16S rRNA. J Microbiol Methods (2013) 95:175-81. doi:10.1016/j.mimet.2013.08.009

28. Mosher JJ, Bowman B, Bernberg EL, Shevchenko O, Kan J, Korlach J, Kaplan L a. Improved performance of the PacBio SMRT technology for 16S rDNA sequencing. J Microbiol Methods (2014) 104:59–60. doi:10.1016/j.mimet.2014.06.012

29. Wagner J, Coupland P, Browne HP, Lawley TD, Francis SC, Parkhill J. Evaluation of PacBio sequencing for full-length bacterial 16S rRNA gene classification. BMC Microbiol (2016) 16:274. doi:10.1186/s12866-016-0891-4

30. Singer E, Bushnell B, Coleman-Derr D, Bowman B, Bowers RM, Levy A, Gies EA, Cheng JF, Copeland A, Klenk H-P, et al. High-resolution phylogenetic microbial community profiling. ISME J (2016) 10:2020–2032. doi:10.1038/ismej.2015.249

31. Ma X, Stachler E, Bibby K. Evaluation of Oxford Nanopore MinION Sequencing for 16S rRNA Microbiome Characterization. bioRxiv (2017). doi:10.1101/099960

32. Benítez-Páez A, Portune KJ, Sanz Y. Species-level resolution of 16S rRNA gene amplicons sequenced through the MinION^TM^ portable nanopore sequencer. Gigascience (2016) 5:4. doi:10.1186/s13742-016-0111-z

33. Benítez-Páez A, Sanz Y. Multi-locus and long amplicon sequencing approach to study microbial diversity at species level using the MinION^TM^ portable nanopore sequencer. Gigascience (2017) doi:10.1093/gigascience/gix043

34. Mitsuhashi S, Kryukov K, Nakagawa S, Takeuchi JS, Shiraishi Y, Asano K, Imanishi T. A portable system for metagenomic analyses using nanopore-based sequencer and laptop computers can realize rapid on-site determination of bacterial compositions. bioRxiv (2017) https://doi.org/10.1101/101865

35. Shin J, Lee S, Go M-J, Lee SY, Kim SC, Lee C-H, Cho B-K. Analysis of the mouse gut microbiome using full-length 16S rRNA amplicon sequencing. Sci Rep (2016) 6:29681. doi:10.1038/srep29681

36. Cuscó A, Sánchez A, Altet L, Ferrer L, Francino O. Individual Signatures Define Canine Skin Microbiota Composition and Variability. Front Vet Sci (2017) 4:6. doi:10.3389/fvets.2017.00006

37. Alm E, Oerther D, Larsen N, Stahl D. The oligonucleotide probe database. Appl Environ Microbiol. (1996) 62:3557–3559

38. Mao DP, Zhou Q, Chen CY, Quan ZX. Coverage evaluation of universal bacterial primers using the metagenomic datasets. BMC Microbiol (2012) 12:66. doi:10.1186/1471-2180-12-66

39. Watson M, Thomson M, Risse J, Talbot R, Santoyo-Lopez J, Gharbi K, Blaxter M. PoRe: An R package for the visualization and analysis of nanopore sequencing data. Bioinformatics (2015) 31:114–115. doi:10.1093/bioinformatics/btu590

40. Wick R. Porechop. Available at: https://github.com/rrwick/Porechop

41. Caporaso JG, Kuczynski J, Stombaugh J, Bittinger K, Bushman FD, Costello EK, Fierer N, Peña AG, Goodrich JK, Gordon JI, et al. QIIME allows analysis of high-throughput community sequencing data. Nat Methods (2010) 7:335–336. doi:10.1038/nmeth.f.303

42. Leggett RM, Heavens D, Caccamo M, Clark MD, Davey RP. NanoOK: multi-reference alignment analysis of nanopore sequencing data, quality and error profiles. Bioinformatics (2015) 32:btv540. doi:10.1093/bioinformatics/btv540

43. Kielbasa SM, Wan R, Sato K, Horton P, Frith MC. Adaptive seeds tame genomic sequence comparison. Genome Res (2011) 21:487–493. doi:10.1101/gr.113985.110

44. DeSantis TZ, Hugenholtz P, Larsen N, Rojas M, Brodie EL, Keller K, Huber T, Dalevi D, Hu P, Andersen GL. Greengenes, a chimera-checked 16S rRNA gene database and workbench compatible with ARB. Appl Environ Microbiol (2006) 72:5069–72. doi:10.1128/AEM.03006-05

45. McDonald D, Price MN, Goodrich J, Nawrocki EP, DeSantis TZ, Probst A, Andersen GL, Knight R, Hugenholtz P. An improved Greengenes taxonomy with explicit ranks for ecological and evolutionary analyses of bacteria and archaea. ISME J (2012) 6:610–8. doi:10.1038/ismej.2011.139

46. Rognes T, Flouri T, Nichols B, Quince C, Mahé F. VSEARCH: a versatile open source tool for metagenomics. (2016)1–22. doi:10.7717/peerj.2584

47. Wang Q, Garrity GM, Tiedje JM, Cole JR. Naive Bayesian classifier for rapid assignment of rRNA sequences into the new bacterial taxonomy. Appl Environ Microbiol (2007) 73:5261–7. doi:10.1128/AEM.00062-07

48. Caporaso JG, Bittinger K, Bushman FD, Desantis TZ, Andersen GL, Knight R. PyNAST: A flexible tool for aligning sequences to a template alignment. Bioinformatics (2010) 26:266–267. doi:10.1093/bioinformatics/btp636

49. Navas-molina JA, Peralta-sánchez JM, González A, McMurdie PJ, Vázquez-Baeza Y, Xu Z, Ursell LK, Lauber C, Zhou H, Song SJ, et al. Advancing Our Understanding of the Human Microbiome Using QIIME. 1st ed. Elsevier Inc. (2013). doi:10.1016/B978-0-12-407863-5.00019-8

50. Fukushima M, Kakinuma K, Kawaguchi R. Phylogenetic analysis of *Salmonella*, *Shigella*, and *Escherichia coli* strains on the basis of the gyrB gene sequence. J Clin Microbiol (2002) 40:2779–85. doi:10.1128/JCM.40.8.2779-2785.2002

51. Lal D, Verma M, Lal R. Exploring internal features of 16S rRNA gene for identification of clinically relevant species of the genus *Streptococcus*. Ann Clin Microbiol Antimicrob (2011) 10:28. doi:10.1186/1476-0711-10-28

52. Oh J, Byrd AL, Deming C, Conlan S, Barnabas B, Blakesley R, Bouffard G, Brooks S, Coleman H, Dekhtyar M, et al. Biogeography and individuality shape function in the human skin metagenome. Nature (2014) 514:59–64. doi:10.1038/nature13786

53. Walters WA, Caporaso JG, Lauber CL, Berg-Lyons D, Fierer N, Knight R. PrimerProspector: de novo design and taxonomic analysis of barcoded polymerase chain reaction primers. Bioinformatics (2011) 27:1159–61. doi:10.1093/bioinformatics/btr087

54. Camanocha A, Dewhirst FE. Host-associated bacterial taxa from Chlorobi, Chloroflexi, GN02, Synergistetes, SR1, TM7, and WPS-2 Phyla/candidate divisions. J Oral Microbiol (2014) 6:25468 doi:10.3402/jom.v6.25468

55. Wallis C, Marshall M, Colyer A, O’Flynn C, Deusch O, Harris S. A longitudinal assessment of changes in bacterial community composition associated with the development of periodontal disease in dogs. Vet Microbiol (2015) 181:271–282. doi:10.1016/j.vetmic.2015.09.003

56. Lin WR, Chen YS, Liu YC. Cellulitis and Bacteremia Caused by *Bergeyella zoohelcum*. J Formos Med Assoc (2007) 106:573–576. doi:10.1016/S0929-6646(07)60008-4

57. Oehler RL, Velez AP, Mizrachi M, Lamarche J, Gompf S. Bite-related and septic syndromes caused by cats and dogs. Lancet Infect Dis (2009) 9:439–447. doi:10.1016/S1473-3099(09)70110-0

58. Sturgeon A, Stull JW, Costa MC, Weese JS. Metagenomic analysis of the canine oral cavity as revealed by high-throughput pyrosequencing of the 16S rRNA gene. Vet Microbiol (2013) 162:891–898. doi:10.1016/j.vetmic.2012.11.018

59. Andersen BM, Steigerwalt AG, O’Connor SP, Hollis DG, Weyant RS, Weaver RE, Brenner DJ. *Neisseria weaveri* sp. nov., formerly CDC group M-5, a gram-negative bacterium associated with dog bite wounds. J Clin Microbiol (1993) 31:2456–66.

60. Kim SY, Adachi Y. Biological and genetic classification of canine intestinal lactic acid bacteria and bifidobacteria. Microbiol Immunol (2007) 51:919–28.

61. Silva BC, Jung LR, Sandes SH, Alvim LB, Bomfim MR, Nicoli JR, Neumann E, Nunes AC. In vitro assessment of functional properties of lactic acid bacteria isolated from faecal microbiota of healthy dogs for potential use as probiotics. Benef Microbes (2013) 4:267–75. doi:10.3920/BM2012.0048

62. Karst SM, Dueholm MS, McIlroy SJ, Kirkegaard RH, Nielsen PH, Albertsen M. Thousands of primer-free, high-quality, full-length SSU rRNA sequences from all domains of life. bioRxiv (2016). doi: https://doi.org/10.1101/070771

63. Links MG, Dumonceaux TJ, Hemmingsen SM, Hill JE. The Chaperonin-60 Universal Target Is a Barcode for Bacteria That Enables De Novo Assembly of Metagenomic Sequence Data. PLoS One (2012) 7: doi:10.1371/journal.pone.0049755

64. Smith AM, Jain M, Mulroney L, Garalde DR, Akeson M. Reading canonical and modified nucleotides in 16S ribosomal RNA using nanopore direct RNA sequencing. bioRxiv (2017) doi: https://doi.org/10.1101/132274

